# Effects of spectral light quality on growth, photosynthetic pigments and bioactive compounds in *Brassicaceae* microgreens

**DOI:** 10.64898/2026.09.01.748622

**Authors:** Valeria González, Gastón Quero, Venancio Riella, Fernanda Zaccari, Ana Cecilia Silveira

## Abstract

LED spectral composition is an important tool for improving the growth and nutritional quality of microgreens cultivated in controlled environments. This study evaluated the effects of three LED light treatments on growth, morphology, pigments, primary metabolites, phenolic composition, and antioxidant capacity in arugula (*Eruca sativa*), mustard (*Brassica juncea*), and radish (*Raphanus sativus*) microgreens. Microgreens were cultivated under controlled environmental conditions and exposed to broad-spectrum white (W), blue-enriched white (WB), and red-enriched white (R) light at a photosynthetic photon flux density of 200 micromol/m2/s. Light quality did not affect yield in any species. However, R increased cotyledon area in arugula by 50 to 60% and promoted hypocotyl elongation in both arugula and radish, whereas W resulted in the longest hypocotyls in mustard. Photosynthetic pigment composition responded differently among species. In mustard, WB increased the chlorophyll a/b ratio (1.12 to 1.18), whereas lutein concentration decreased from 7.06 to 4.20 mg 100 g/FW. Primary metabolism also responded to light treatments in a species-dependent manner. In mustard, W increased glucose (0.43 vs. 0.26 and 0.29 g 100 g/ FW) and fructose (0.33 vs. 0.20 and 0.22 g 100 g/ FW) concentrations compared with WB and R. Organic acid composition was more responsive to light treatments in radish, with higher concentrations under R. Phenolic metabolism also responded in a species-dependent manner. In mustard, W increased total phenolic content to 0.25 mg GAE g/FW compared with 0.15 mg GAE g/FW under WB and R, and ABTS antioxidant capacity to 1.17 mg TE g/FW compared with 0.74 and 0.75 mg TE g/FW under WB and R, respectively. Individual phenolic compounds were also affected by light treatments, particularly in arugula and mustard. These findings demonstrate that the effects of LED spectral composition on microgreen quality are highly species-dependent. Therefore, LED light spectra should be optimized according to the target species and the desired quality attributes rather than applying a single lighting strategy to all Brassicaceae microgreens.

## Introduction

Microgreens are young seedlings with expanded cotyledons and their first pair of true leaves (Zhang et al., 2021). They are harvested 10–20 days post-emergence, at which point they reach approximately 5–10 cm in height, depending on the species (Kyriacou et al., 2016; Flores et al., 2022). These products have attracted interest due to their high nutritional value and bioactive compound content, which in many cases is higher than that of adult plants (Alloggia et al., 2023; Drozdowska et al., 2020; Rai et al., 2022). In particular, species from the Brassicaceae family are among the most widely used, mainly due to their ease of cultivation, rapid growth, and composition (Xiao et al., 2019; Li et al., 2023a).

Microgreens are generally cultivated in controlled environment systems, where lighting management is a critical factor because it is one of the main environmental regulators of plant growth and development (Farhangi et al., 2025; Ying et al., 2020a). Since lighting can be fully modulated in controlled environments, spectral quality becomes a key tool for directing morphophysiological responses in plants. In this regard, LED (Light-Emitting Diode) lighting offers advantages over traditional lamps due to its lower energy consumption, lower heat output, longer lifespan, and the ability to adjust spectral quality (Shibaeva et al., 2022).

Within the visible spectrum, the wavelengths most absorbed by pigments and photoreceptors are blue (400–500 nm) and red (600–700 nm) light (Mlinarić et al., 2023; Zhang et al., 2020). As a result, most lighting recipes used in indoor cultivation are dichromatic (Cowden et al., 2024; Ying et al., 2020b). Some studies have reported that blue light regulates stomatal opening, the accumulation of antioxidant compounds, and reduces hypocotyl elongation (Thongtip et al., 2024; Toscano et al., 2021). Conversely, red light promotes photosynthesis, height growth, and the accumulation of fresh and dry biomass (Liang et al., 2022; Orlando et al., 2022). However, several studies have shown that the red-to-blue (R: B) ratio can have a more decisive effect than its individual components, modulating morphophysiological responses differentially across species (Haghighi et al., 2025). Additionally, recent studies suggest that broad-spectrum lighting can promote more favorable growth and chemical composition responses than spectra composed of one or two wavelengths (Farhangi et al., 2025; Orlando et al., 2022). Others have reported that combining white light with varying proportions of red and blue produces a greater positive effect (Jasenovska et al., 2024). However, the responses of microgreens species to spectral composition remain species-dependent and are not yet fully understood. Therefore, the objective of this study was to investigate how different spectral compositions influence morphological and biochemical traits in microgreens of three *Brassicaceae* species.

## Material and methods

### Plant material, growth conditions and light treatments

Three species from the Brassicaceae family were used: arugula (*Eruca sativa*) variety Forte (Profit Seed, Italy), mustard (*Brassica juncea*) variety Green Boy (Takii Seed, Japan), and radish (*Raphanus sativus*) variety Saxa (Sais, Italy). Sowing was carried out by broadcasting in polypropylene trays (52 x 25.5 x 3.5 cm) on a substrate composed of compost, perlite, and vermiculite (3:0.5:0.5 v/v/v, Terrafértil, Argentina) previously sterilized (121 °C for 20 min). A sowing density of 2.5 plants/cm ² was used for arugula and mustard, and 1.5 plants/cm ² for radish.

After sowing, the trays were randomly assigned to one of the three light treatments and placed on the corresponding shelves in a controlled room where temperature and relative humidity were kept constant (20 °C; 76% RH on average). The shelves were covered with an opaque material to prevent light pollution, and each shelf had a configurable LED grow light bar (Smart foldable LED grow light bar SFB-200 W, China) for applying light treatments. Immediately after sowing, the trays were kept in the dark for 4 days to promote germination. Three light treatments differing in spectral composition were established: broad-spectrum white light (W), blue-enriched white light (WB), and red-enriched white light (R). A light intensity of 200 µmol/m²/s and a photoperiod of 12 h were used. The spectral characteristics of the three LED treatments, including spectral irradiance, integrated irradiance (Ee), PPFD, and the red-to-blue ratio (R:B), are presented in Figure 1. The spectral characteristics and PPFD of each treatment were determined using a UPRtek MK350S LED meter (Miaoli County, Taiwan).

**Fig. 1.**
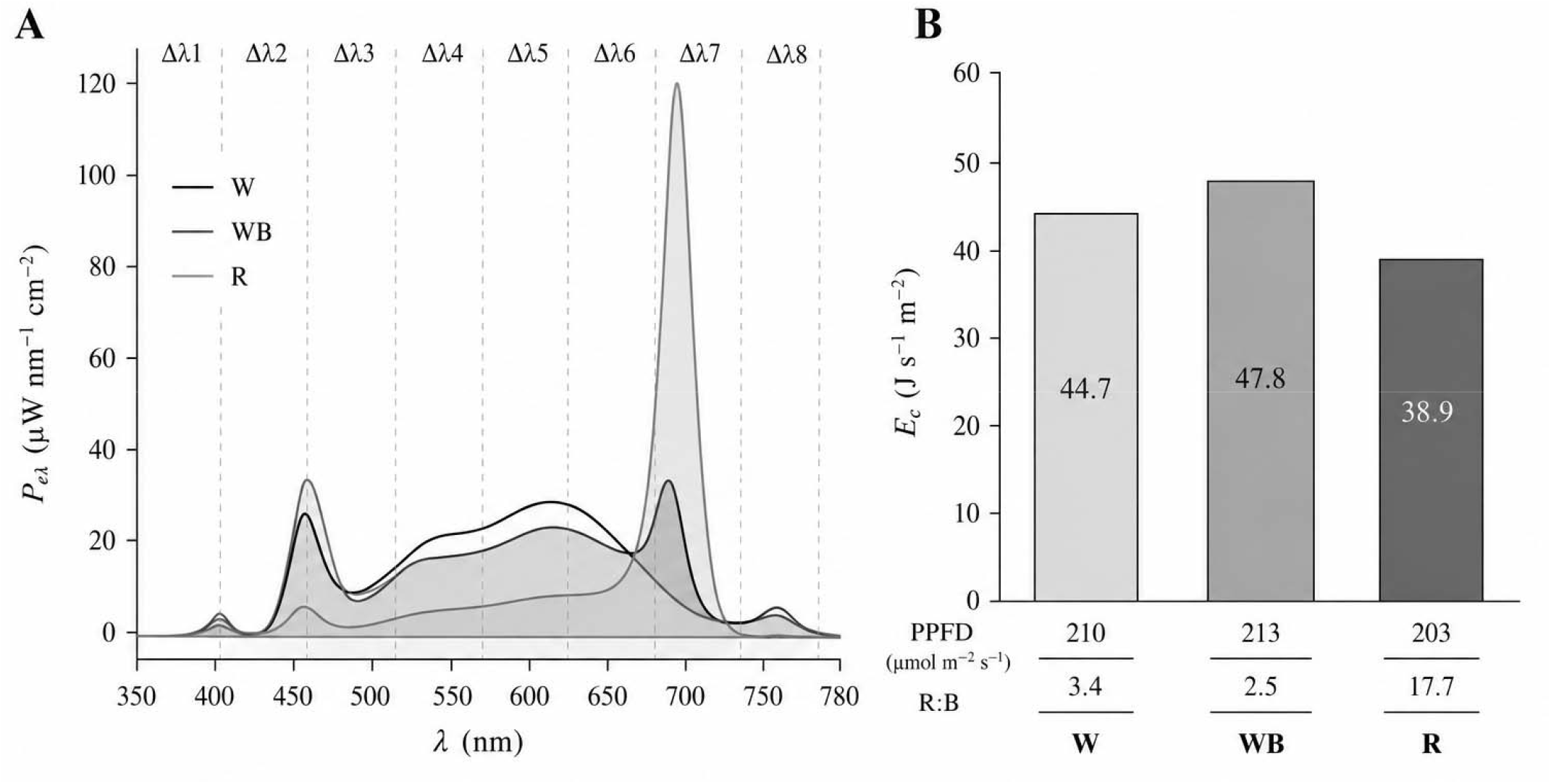
Spectral characterization of the LED light treatments. (A) Spectral irradiance (P eλ) over the 350–780 nm wavelength range. Dashed vertical lines indicate the wavebands (Δλ1–Δλ8) used for the analysis. (B) Integrated irradiance (Ee) within the photosynthetically active radiation (PAR, 400–700 nm) range. Photosynthetic photon flux density (PPFD) and the red-to-blue ratio (R:B) are indicated below each bar. W, white light; WB, blue-enriched white light; R, red-enriched light.

Harvesting was done manually with scissors, making a cut flush with the substrate, 17 days after sowing for arugula and mustard, and 10 days after sowing for radish, when the plants had expanded cotyledons and their first pair of true leaves.

### Biomass yield

To determine yield, the entire harvest from each tray was weighed on a digital scale (Ohaus, Scout™ Pro SP602, USA). Yield was expressed as grams of fresh weight per square meter (g·m ²).

### Dry matter of roots and shoots

To determine the percentage of dry matter corresponding to the roots and shoots at harvest, three 9.5 cm² portions were taken from each tray, from which roots and shoots were separated. Each fraction was dried in a domestic microwave oven (Enxuta, Brazil, 1150 W, 20 L) at 800 W for 6 minutes for roots and 12 minutes for the shoots. The result was expressed as a percentage of dry matter.

### Hypocotyl length, cotyledon area, and elongation rate

Ten seedlings per replicate were taken 7 days after sowing and at harvest. Hypocotyl length and cotyledon area were determined at these times using ImageJ software (version 1.42, National Institutes of Health, USA). Based on this, the daily hypocotyl elongation rate (cm·day ¹) was calculated using the methodology proposed by Kong et al. (2019).

### Color

Color was determined on 20 cotyledons per replicate using a colorimeter (PCE-TCR 200, Spain) based on the CIE Lab color space. Measurements were taken on the adaxial surface at the center of each cotyledon against a white background. Because of the small size of the cotyledons, a custom paper positioning template was used to ensure consistent placement during color measurements. The template consisted of a circular guide matching the diameter of the colorimeter measuring aperture and alignment marks that allowed each cotyledon to be centered over the measuring aperture. The parameters luminosity (L*), a*, and b* were recorded, from which chroma (C*) and Hue angle (h_ab_) were calculated.

### Chlorophyll and Total Carotenoids

For extraction, 0.3 g of plant material was homogenized in 10 mL of acetone: water (80:20 v/v) and 1 g/L butylhydroxytoluene at 28,000 rpm for 30 s (Scientz XHF-D, China). The homogenate was kept in the dark for 1 h at 7 °C and then centrifuged at 10,500 rpm for 10 min at 4 °C. The supernatant was then separated, and the absorbance was measured at 663, 646, and 470 nm to determine the content of chlorophyll a, chlorophyll b, and total carotenoids, respectively. The determination was performed using a UV-Vis spectrophotometer (Genesys, Thermo Scientific, USA). Chlorophyll a, chlorophyll b, and total carotenoids were calculated according to Lichtenthaler and Wellburn (1983) using the following equations:

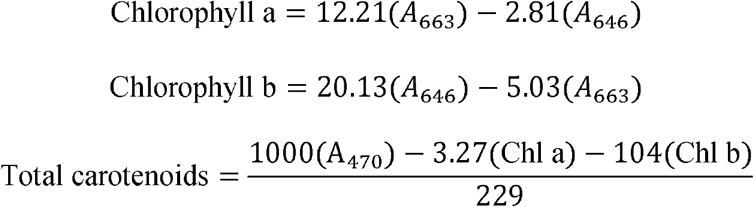

### Chlorophyll and lutein determination by HPLC-DAD

The same extract used for the spectrophotometric determination was used for HPLC analysis. The extract was filtered through 0.45 µm PTFE membrane filters and analyzed using the HPLC system described above. Separation was performed on a YMC Carotenoid C30 column (250 × 4.6 mm i.d., 5 µm; YMC Co., Ltd., Kyoto, Japan) maintained at 30 °C. Isocratic elution was carried out using ethanol:methanol:tetrahydrofuran (75:20:5, v/v/v) as the mobile phase at a flow rate of 0.7 mL min ¹. Chlorophyll a was quantified at 665 nm, whereas chlorophyll b and lutein were quantified at 450 nm. Pigments were identified by comparing their retention times and UV–visible absorption spectra with those of authentic standards (Sigma-Aldrich, St. Louis, MO, USA) and quantified using external calibration curves. Results were expressed as mg 100 g ¹ fresh weight (FW).

### Extraction for TAC and TP determinations

For the extraction, 0.5 g of plant material was weighed, and 3 mL of methanol: water (70:30 v/v) was added. The mixture was then homogenized (Scientz, XHF-D, China) at 28,000 rpm for 30 s, followed by centrifugation (15,000 × g, 4 °C, Thermo Scientific, Sorvall™ ST 16R, Germany) for 10 min. The supernatant was used for the determinations.

### Total Antioxidant Capacity measurement

Total antioxidant capacity (TAC) was determined spectrophotometrically (Multiskan Sky, Thermo Scientific, USA) using the DPPH, FRAP, and ABTS methods. The DPPH method was performed according to Brand-Williams et al. (1995), with modifications. For the reaction, 10 µL of extract and 190 µL of DPPH solution with absorbance adjusted to 1.1 were used. The samples were incubated in the dark for 105 min (arugula and mustard) and 90 min (radish), and the absorbance was subsequently measured at 515 nm.

The FRAP reagent was prepared with a 300 mM sodium acetate (C H NaO) solution at pH 3.6, 10 mM 2,4,6-tripyridyl-s-triazine solution with 40 mM hydrochloric acid (HCl), and a 20 mM ferric chloride (FeCl) solution, in a 10:1:1 v/v/v ratio, previously incubated at 37 °C for 30 min (Benzie and Strain, 1999). For the determination, 10 µL of the extract and 190 µL of FRAP reagent were used. The absorbance was determined at 593 nm.

Finally, the ABTS method was performed as proposed by Re et al. (1999), using 10 µL of the extract to which 190 µL of ABTS solution were added, with absorbance adjusted to 0.9. Incubation times were 75, 60, and 120 min for arugula, radish, and mustard, respectively. Absorbance was determined at 734 nm.

In all cases, the results were expressed as mg Trolox equivalents (Merck KGaA, Darmstadt, Germany) per gram of fresh weight (mg TE g ¹ FW).

### Total Polyphenols determination

For the determination, 30 µL of 1 N Folin-Ciocalteu reagent, 200 µL of a 0.4% sodium hydroxide (NaOH) and 2% sodium carbonate (Na CO) solution, and 20 µL of the extract were used, according to the methodology proposed by Singleton and Rossi (1965), with modifications. The samples were incubated for 60, 30, and 75 min for arugula, mustard, and radish, respectively. Absorbance was determined at 765 nm, and the results were expressed as milligrams of gallic acid equivalents (Merck KGaA, Darmstadt, Germany) per gram of fresh weight (mg AGE g^-1^ WF).

### Individual polyphenols determination

For the quantification of individual polyphenols, the same extract obtained for the determination of PT and TAC was used, filtered through 0.45 µm PTFE membrane filters (Merck Millipore, Darmstadt, Germany). The samples were injected into an HPLC-DAD system equipped with an autosampler (SIL-20AC; Shimadzu Corporation, Kyoto, Japan) and a diode array detector (DAD-20A). A Restek C18 column (5 µm, 250 × 6 mm; Restek, Bellefonte, Pennsylvania, USA) was used. The mobile phase consisted of 1 % formic acid in water (A) and 100 % acetonitrile (B), using a gradient elution program as follows: 0 min, 5 % B; 20 min, 50 % B; 25 min, 60 % B; 29 min, 60 % B; 30 min, 5 % B; and 35 min, 0 % B. The flow rate was 0.9 mL/min. The column oven was maintained at 35 °C, the run time was 35 min, and the injection volume was 10 µL.

Identification was performed by comparison of retention times and absorption spectra with commercial standards. Quantification was carried out at 254, 280, 320, 360 and 520 nm according to the phenolic family. Calibration curves were prepared using commercial standards. The results were expressed as mg per gram of fresh weight (mg AGE g^-1^ WF).

### Determination of Sugars and Organic Acids

For extraction, 1 g of plant material was homogenized with 10 mL of distilled water using a homogenizer (Scientz, XHF-D, China) at 28,000 rpm for 60 s. The homogenate was centrifuged at 15,000 × g for 20 min at 4 °C, and the supernatant was recovered and stored at −20 °C until analysis. Prior to injection, the extract was filtered through 0.45 µm PTFE membrane filters (Merck Millipore, Darmstadt, Germany) and injected into an HPLC system (SIL-20AC autosampler; Shimadzu Corporation, Kyoto, Japan) equipped with a refractive index detector (RID-10A) and a diode array detector (DAD-20A).

For sugars (glucose, fructose, and sucrose), separation was performed on a Luna® Omega 3 µm SUGAR 100 Å column (250 × 4.6 mm; Phenomenex, Torrance, CA, USA) using an acetonitrile: water mobile phase (80:20 v/v) in isocratic mode. Detection was performed using refractive index (RI). For organic acids (citric, malic, and ascorbic), determination was performed by DAD at 210 and 254 nm using a Shodex KC-811 RSpak column (8 x 300 mm) with 0.1% phosphoric acid in water mobile phase in isocratic mode. For both determinations, a flow rate of 1 mL/min, a column temperature of 40 °C, and a run time of 20 min were used. Identification was performed by comparison of retention times with commercial standards, and quantification was carried out using calibration curves prepared with commercial standards. Results were expressed in milligrams per gram of fresh weight (mg g^-1^ FW).

### Experimental design and data analysis

For each species, the experiment was conducted using a completely randomized design, with light treatment as the single factor and four independent replicates per treatment. Each replicate consisted of one cultivation tray placed on an individually illuminated shelf that was optically isolated from the remaining experimental units. Light treatments were randomly assigned to the shelves. Data were analyzed by one-way analysis of variance (ANOVA). When treatment effects were significant, means were compared using Tukey’s honestly significant difference (HSD) test at P < 0.05. Statistical analyses were performed using R software (R Core Team, 2026) within the RStudio integrated development environment (Posit Team, 2023). Additionally, a heatmap was generated using standardized values (z-scores) to facilitate visualization of treatment responses within each species.

## Results and discussion

### Biomass yield, dry matter content and morphological traits

No significant differences in yield were detected among light treatments in any of the three species. However, shoot dry matter content was affected by light quality. In arugula, treatment WB resulted in the highest values, approximately twofold higher than those observed under W and R, which did not differ from each other. In mustard, treatment W showed higher shoot dry matter content than WB, whereas R presented intermediate values and did not differ significantly from either treatment. In radish, treatment R showed the highest values, being approximately 40% higher than W, while WB showed intermediate values.

At the root level, no significant differences were detected between treatments in mustard or radish, with average values of 9.01% and 13.85%, respectively. In arugula, treatment R showed the highest dry matter content, while W presented intermediate values and did not differ significantly from either treatment.

Light quality significantly affected hypocotyl length in all three species. In arugula, seedlings grown under treatment R developed longer hypocotyls than those exposed to treatment W. A similar pattern was observed in radish, where hypocotyl length ranged from 4.69 to 6.15 cm, with the longest hypocotyls recorded under R and the shortest under W. In contrast, mustard responded differently to light quality, as seedlings grown under treatment W exhibited the greatest hypocotyl length, whereas WB resulted in the shortest values and R showed intermediate values (Table 1).

**Table 1.** Biomass yield, dry matter content, and morphological traits of arugula, mustard, and radish microgreens grown under different LED light treatments.

| Light treatment | Arugula | Mustard | Radish |
| --- | --- | --- | --- |
| **\*Fresh biomass yield (g m<sup>-2</sup>)** |  |  |  |
| **W** | 673.60 ± 46.19 a | 969.13 ± 43.76 a | 1,593.90 ± 59.45 a |
| **WB** | 470.25 ± 38.52 a | 948.18 ± 29.69 a | 1,438.55 ± 68.86 a |
| **R** | 508.50 ± 34.64 a | 957.23 ± 33.05 a | 1,652.33 ± 23.02 a |
| **\*\*Dry matter of shoots (%)** |  |  |  |
| **W** | 6.05 ± 0.75 b | 6.01 ± 0.18 a | 4.98 ± 0.36 b |
| **WB** | 10.23 ± 1.04 a | 3.40 ± 0.33 b | 6.38 ± 0.24 ab |
| **R** | 3.83 ± 0.28 b | 4.97 ± 0.73 ab | 7.09 ± 0.63 a |
| **\*\*Dry matter of roots (%)** |  |  |  |
| **W** | 7.45 ± 1.31 ab | 10.43 ± 0.91 a | 10.18 ± 2.27 a |
| **WB** | 3.66 ± 0.54 b | 6.84 ± 2.25 a | 16.95 ± 1.29 a |
| **R** | 10.04 ± 1.83 a | 9.77 ± 0.77 a | 14.43 ± 1.83 a |
| **\*\*\*Hypocotyl length (cm)** |  |  |  |
| **W** | 2.24 ± 0.07 b | 4.35 ± 0.14 a | 4.69 ± 0.30 b |
| **WB** | 2.42 ± 0.12 ab | 3.45 ± 0.15 b | 5.30 ± 0.25 ab |
| **R** | 2.62 ± 0.06 a | 3.95 ± 0.13 ab | 6.15 ± 0.12 a |
| **\*\*\*Hypocotyl elongation rate (cm d<sup>-1</sup>)** |  |  |  |
| **W** | 0.09 ± 0.01 ab | 0.20 ± 0.01 a | 0.29 ± 0.08 a |
| **WB** | 0.14 ± 0.02 a | 0.10 ± 0.01 b | 0.23 ± 0.04 a |
| **R** | 0.06 ± 0.01 b | 0.11 ± 0.01 b | 0.30 ± 0.05 a |
| **\*\*\*Cotyledon area (cm<sup>2</sup>)** |  |  |  |
| **W** | 0.78 ± 0.06 b | 0.38 ± 0.03 a | 1.33 ± 0.08 a |
| **WB** | 0.82 ± 0.03 b | 0.38 ± 0.02 a | 1.04 ± 0.06 a |
| **R** | 1.24 ± 0.04 a | 0.35 ± 0.03 a | 1.38 ± 0.05 a |

Values are expressed as mean ± standard error (\*n = 4, \*\*n = 12, \*\*\*n = 10). Different lowercase letters within each species indicate significant differences among light treatments according to Tukey's test (P < 0.05).

In arugula, the highest elongation rate was observed under WB, whereas R showed the lowest rate (0.06 cm day ¹), while W presented intermediate values and did not differ significantly from either treatment. In mustard, treatment W resulted in the highest elongation rate (0.20 cm day ¹), whereas WB and R showed lower values and did not differ significantly from each other. In radish, no significant differences in elongation rate were detected among light treatments.

Cotyledon area was not affected by the light treatments in radish or mustard. In contrast, it was significantly influenced by light quality in arugula, with treatment R producing values approximately 50–60% higher than those observed under W and WB.

### Color measurement

The color parameters L*, h_ab_, and C* showed different responses depending on the species and the light treatment under which they were grown (Fig. 2). Specifically, in radish, no significant differences were observed in the L*, h_ab_, and C* variables across the three light treatments. In mustard, lightness was affected by the light treatment, with higher L* values found under treatments W and R (66.03 and 66.04, respectively) compared to treatment WB (64.30). Hue angle (h_ab_), it showed a lower value in treatment W (113.17), indicating a more yellow coloration in this treatment. No significant differences were observed for C*. In arugula, treatment R showed greater lightness and chroma than W and WB, whereas hue angle was higher under W and WB. These results indicate that treatment R produced a brighter, more saturated, and less green (or more yellow) appearance than W and WB.

**Fig. 2.**
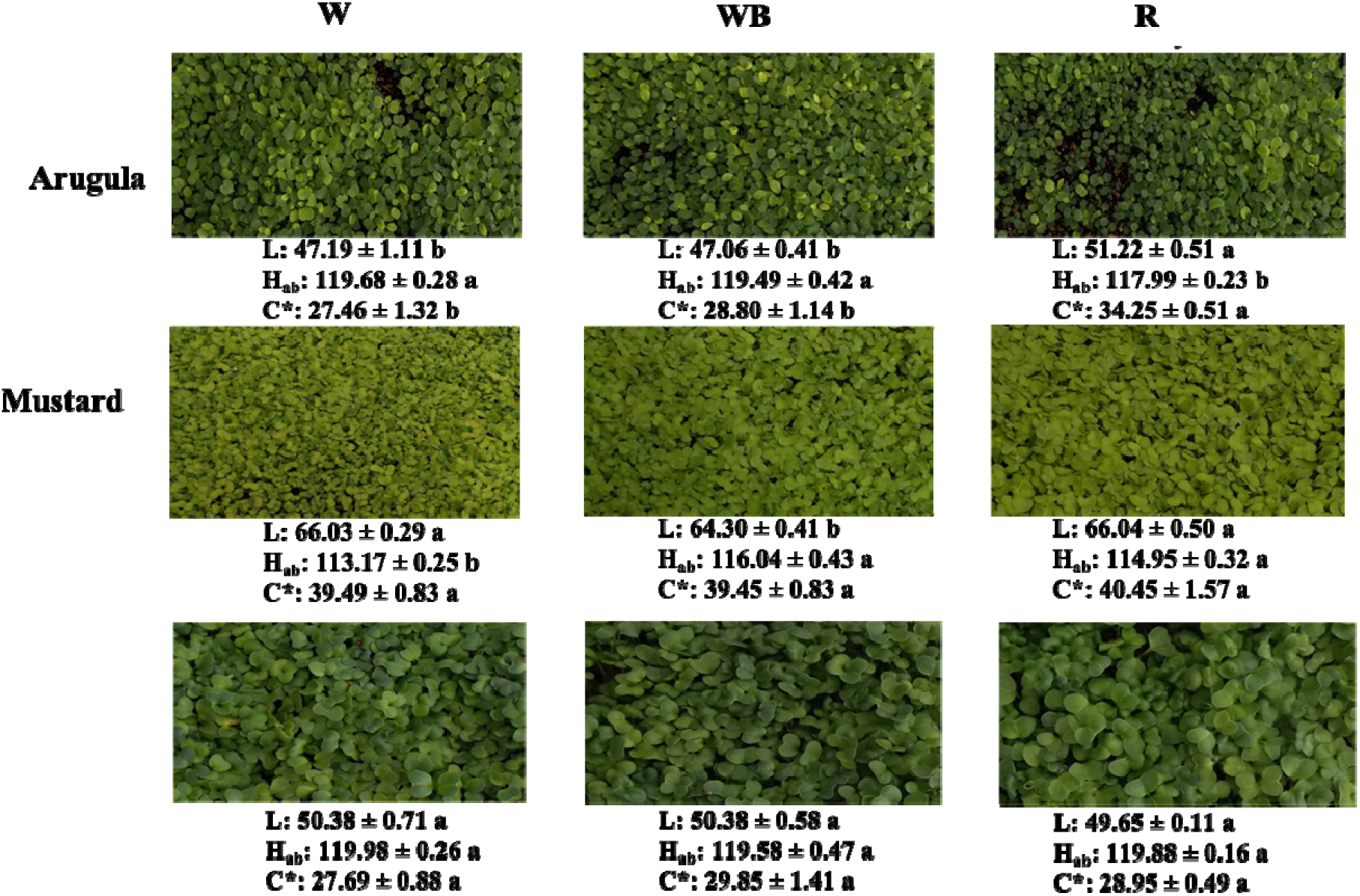
Representative images of arugula, mustard, and radish microgreens at harvest grown under white (W), blue-enriched white (WB), and red-enriched (R) LED light treatments.

### Chlorophylls and Total Carotenoids

Neither chlorophyll a (Chl a) nor chlorophyll b (Chl b) content was altered by light treatments in any of the three species studied (data not shown). In arugula, Chl a content ranged from 0.45 to 0.65 mg g^-1^ FW, while Chl b content ranged from 0.09 to 0.14 mg g^-1^ FW. In mustard, Chl a content ranged from 0.24 to 0.27 mg g^-1^ FW. Chl b content in this species ranged from 0.07 to 0.09 mg g^-1^ FW with no differences between treatments. In radish, Chl a content ranged from 0.35 to 0.42 mg g^-1^ FW, whereas the Chl b content ranged from 0.09 to 0.10 mg g^-1^ FW.

Regarding total carotenoid content, no significant effects of light treatment were observed in any of the three species evaluated (data not shown). In arugula, total carotenoid content ranged from 0.12 to 0.17 mg g^-1^ FW. In mustard, total carotenoid content averaged 0.06 mg g^-1^ FW, whereas in radish, total carotenoid content ranged from 0.08 to 0.10 mg g ¹ FW.

### Individual chlorophylls and lutein

Light treatments affected the composition of photosynthetic pigments differently depending on the species (Figure 3). In mustard, the chlorophyll a/b ratio and lutein concentration was affected by the treatments. The chlorophyll a/b ratio was higher under treatment WB than under W, whereas R showed intermediate values. In contrast, lutein concentration was highest under W, lowest under WB, and intermediate under R. Chlorophyll a, chlorophyll b, and total chlorophyll contents were not affected. In radish, only the chlorophyll a/b ratio was influenced by treatments, with treatment R showing higher values than WB, whereas W was intermediate. No significant differences were observed for chlorophyll a, chlorophyll b, total chlorophyll, or lutein. Likewise, none of the HPLC-determined pigments were affected in arugula.

**Fig. 3.**
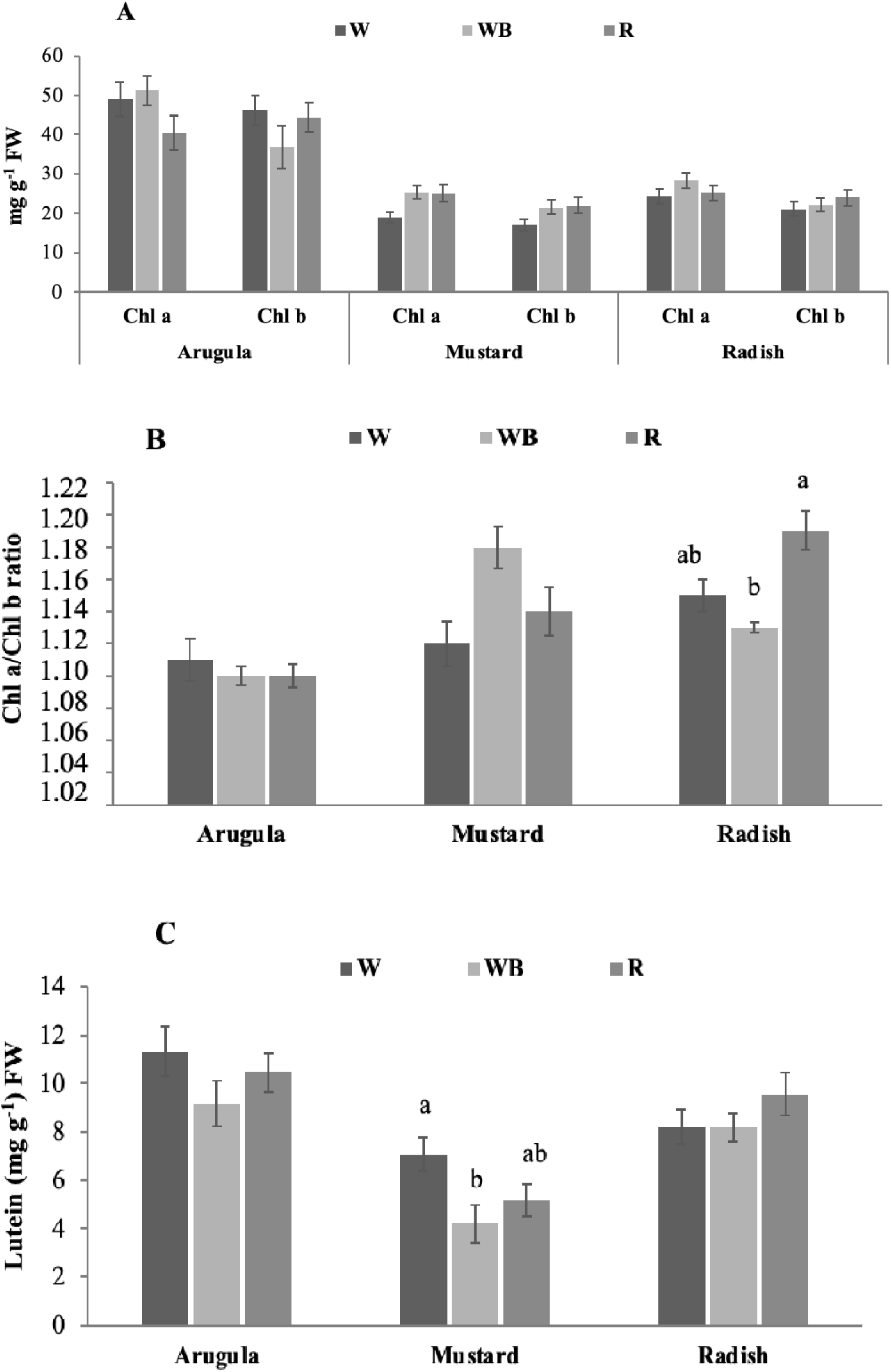
Chlorophyll a, chlorophyll b, chlorophyll a/b ratio, and lutein content in arugula, mustard, and radish microgreens grown under different LED light treatments. Values are expressed as mean ± standard error (n = 4). Different lowercase letters within each species indicate significant differences among light treatments according to Tukey’s test (P < 0.05).

### Sugar and organic acid

The concentrations of glucose and fructose were affected differently by the light treatments depending on the species (Table 2). In arugula, glucose and fructose content were not affected by the light treatments. Fructose was the predominant sugar, with concentrations approximately 2.6 times higher than those of glucose. In radish, no differences were observed between treatments; however, glucose was the major sugar, with concentrations approximately 7 times higher than those of fructose. In mustard, treatment W resulted in higher glucose and fructose content than treatments WB and R. Glucose and fructose were present at similar concentrations in mustard.

**Table 2.** Glucose and fructose concentrations in arugula, mustard, and radish microgreens grown under different LED light treatments.

| Light treatment | Arugula | Mustard | Radish |
| --- | --- | --- | --- |
| **Glucose (g 100 g<sup>-1</sup> FW)** |  |  |  |
| **W** | 0.38 ± 0.14 a | 0.43 ± 0.03 a | 0.41 ± 0.04 a |
| **WB** | 0.42 ± 0.05 a | 0.26 ± 0.02 b | 0.42 ± 0.06 a |
| **R** | 0.49 ± 0.03 a | 0.29 ± 0.02 b | 0.40 ± 0.03 a |
| **Fructose (g 100 g<sup>-1</sup> FW)** |  |  |  |
| **W** | 1.22 ± 0.19 a | 0.33 ± 0.03 a | 0.050 ± 0.003 a |
| **WB** | 0.92 ± 0.22 a | 0.20 ± 0.02 b | 0.060 ± 0.006 a |
| **R** | 1.29 ± 0.12 a | 0.22 ± 0.02 b | 0.050 ± 0.004 a |

Values are expressed as mean ± standard error (n = 4). Different lowercase letters within each species indicate significant differences among light treatments according to Tukey's test (P < 0.05).

In arugula, light quality significantly affected succinic and malic acid concentrations (Table 3). Succinic acid increased under the R treatment compared with W and WB, whereas malic acid decreased under R relative to W, with WB showing intermediate values. No significant differences among light treatments were observed for citric or fumaric acids, and ascorbic acid was not detected.

**Table 3.**
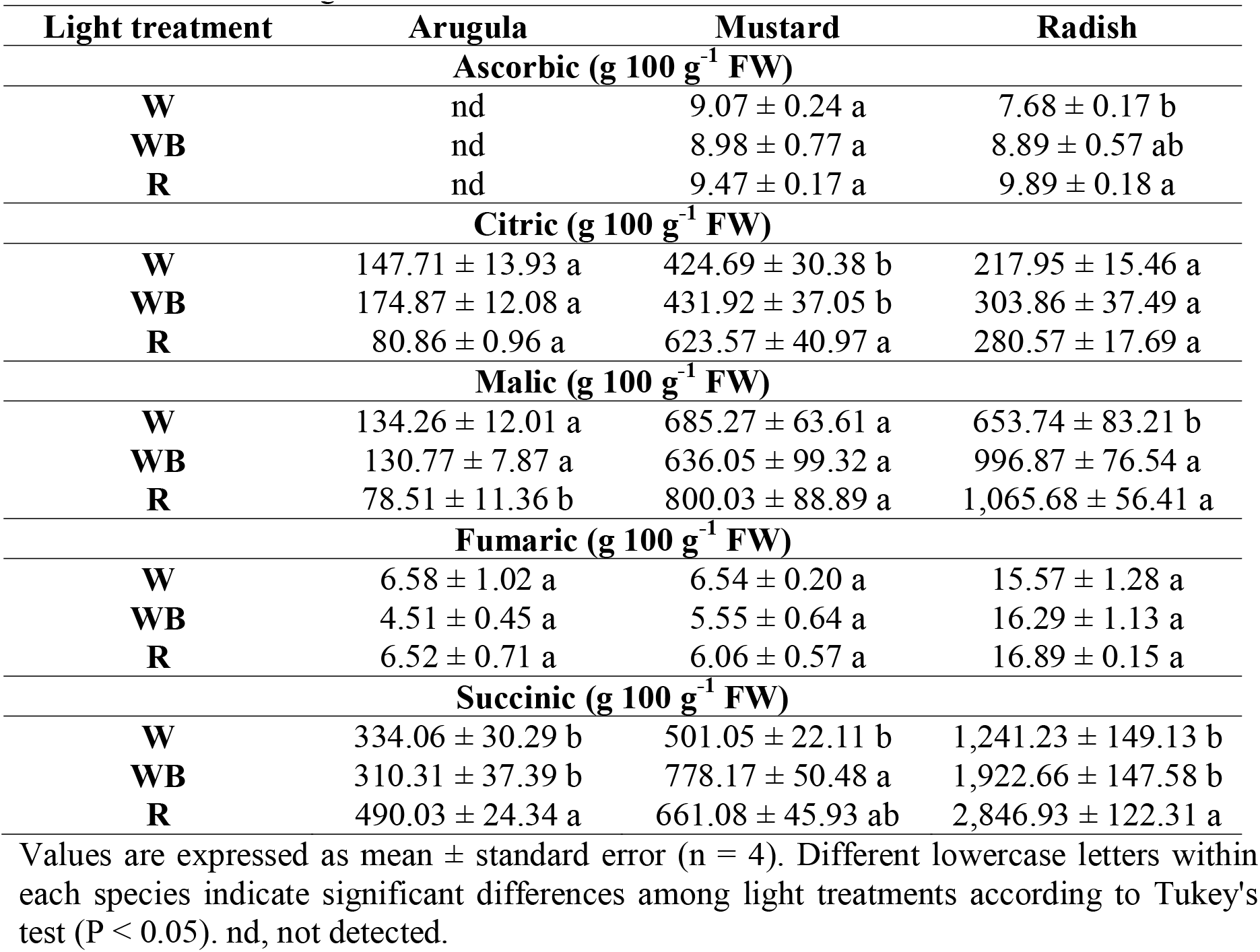
Organic acid concentrations in arugula, mustard, and radish microgreens grown under different LED light treatments.

| Light treatment | Arugula | Mustard | Radish |
| --- | --- | --- | --- |
| **Ascorbic (g 100 g<sup>-1</sup> FW)** |  |  |  |
| **W** | nd | 9.07 ± 0.24 a | 7.68 ± 0.17 b |
| **WB** | nd | 8.98 ± 0.77 a | 8.89 ± 0.57 ab |
| **R** | nd | 9.47 ± 0.17 a | 9.89 ± 0.18 a |
| **Citric (g 100 g<sup>-1</sup> FW)** |  |  |  |
| **W** | 147.71 ± 13.93 a | 424.69 ± 30.38 b | 217.95 ± 15.46 a |
| **WB** | 174.87 ± 12.08 a | 431.92 ± 37.05 b | 303.86 ± 37.49 a |
| **R** | 80.86 ± 0.96 a | 623.57 ± 40.97 a | 280.57 ± 17.69 a |
| **Malic (g 100 g<sup>-1</sup> FW)** |  |  |  |
| **W** | 134.26 ± 12.01 a | 685.27 ± 63.61 a | 653.74 ± 83.21 b |
| **WB** | 130.77 ± 7.87 a | 636.05 ± 99.32 a | 996.87 ± 76.54 a |
| **R** | 78.51 ± 11.36 b | 800.03 ± 88.89 a | 1,065.68 ± 56.41 a |
| **Fumaric (g 100 g<sup>-1</sup> FW)** |  |  |  |
| **W** | 6.58 ± 1.02 a | 6.54 ± 0.20 a | 15.57 ± 1.28 a |
| **WB** | 4.51 ± 0.45 a | 5.55 ± 0.64 a | 16.29 ± 1.13 a |
| **R** | 6.52 ± 0.71 a | 6.06 ± 0.57 a | 16.89 ± 0.15 a |
| **Succinic (g 100 g<sup>-1</sup> FW)** |  |  |  |
| **W** | 334.06 ± 30.29 b | 501.05 ± 22.11 b | 1,241.23 ± 149.13 b |
| **WB** | 310.31 ± 37.39 b | 778.17 ± 50.48 a | 1,922.66 ± 147.58 b |
| **R** | 490.03 ± 24.34 a | 661.08 ± 45.93 ab | 2,846.93 ± 122.31 a |

Values are expressed as mean ± standard error (n = 4). Different lowercase letters within each species indicate significant differences among light treatments according to Tukey's test (P < 0.05). nd, not detected.

In mustard, light quality significantly affected citric and succinic acid concentrations. Citric acid reached its highest concentration under the R treatment, whereas succinic acid was higher under WB than under W, with R showing intermediate values. Malic, fumaric, and ascorbic acid concentrations were not affected by light treatment.

In radish, light quality significantly affected succinic, malic, and ascorbic acid concentrations. Succinic acid reached its highest concentration under the R treatment, approximately 50% higher than under W and WB. Likewise, malic acid concentrations were higher under WB and R than under W. Ascorbic acid concentration was also higher under R than under W, whereas citric and fumaric acids were not significantly affected by light treatment.

### Total polyphenol (TPC) content and antioxidant capacity (TAC)

Total phenolic content and antioxidant capacity showed species-dependent responses to light treatment (Figure 4). Phenolic concentrations ranged from 0.45 to 0.76 mg GAE g ¹ FW. In mustard, values under treatment W resulted in a 37% higher than those under WB and R, which did not differ from each other. In radish, TPC was not affected by the applied light treatment, with an average value of 0.73 mg GAE g^-1^ FW. The same behavior was observed in arugula, where, although the values were slightly lower (0.55-0.65 mg GAE g^-1^ FW), there was no difference under the spectra used.

**Fig. 4.**
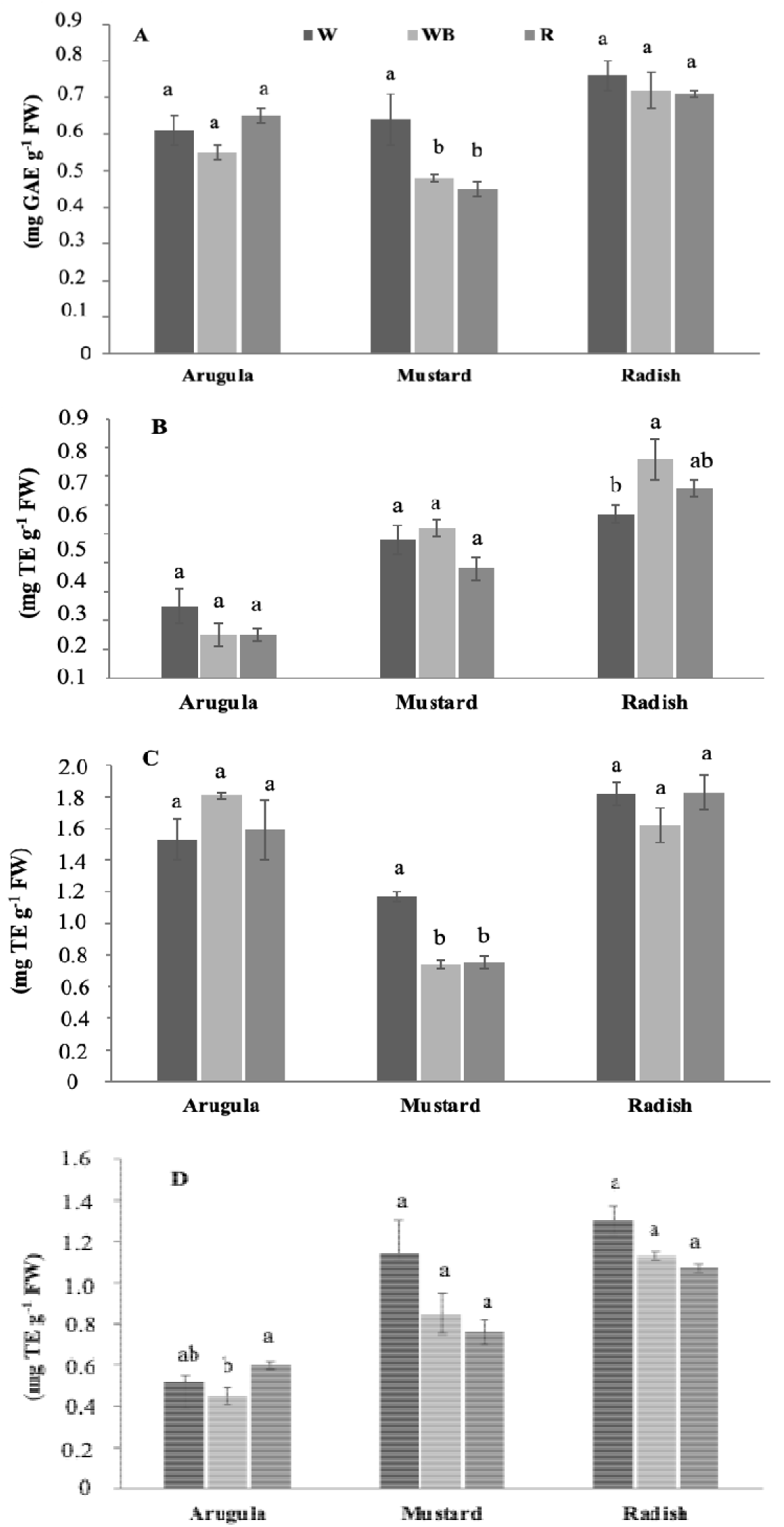
Total phenolic content and antioxidant capacity of arugula, mustard, and radish microgreens grown under different LED light treatments. (A) Total phenolic content (TPC), (B) DPPH radical scavenging activity, (C) ABTS radical scavenging activity, and (D) ferric reducing antioxidant power (FRAP). Values are expressed as mean ± standard error (n = 4). Different lowercase letters within each species indicate significant differences among light treatments according to Tukey’s test (P < 0.05).

In arugula, TAC determined using the DPPH method showed no significant differences among treatments, with an average value of 0.18 mg TE g ¹ FW. Similar results were obtained with the ABTS assay. However, FRAP values were lower under treatment WB than under treatment R, while treatment W showed intermediate values and did not differ from either treatment.

In mustard, TAC measured by DPPH and FRAP showed no significant differences among the treatments, with average values of 0.46 and 0.91 mg TE g^-1^ FW, respectively. In contrast, TAC determined by ABTS was significantly higher under treatment W, with a 57% increase compared to treatments WB and R. In radish, TAC measured by the DPPH method showed differences between treatments, with higher values recorded under treatment WB than under treatment W, while treatment R showed intermediate values and did not differ from either treatment. However, no differences were observed among light spectra when TAC was determined using ABTS or FRAP.

### Individual polyphenols

The individual polyphenols detected in each species under the treatments used are shown in Table 4. In arugula, nine phenolic compounds were identified, two of which were affected by light treatment. In particular, spectral quality influenced the accumulation of 3,4- dihydroxybenzoic acid and gallic acid. In both cases, treatment W resulted in the highest concentrations of these acids, with treatment WB being 88% and 52% lower, respectively. Unlike arugula, mustard microgreens showed greater sensitivity to light quality. In this regard, treatment W promoted greater accumulation of most of the compounds detected. 3,4-dihydroxybenzoic, 4-hydroxybenzoic, chlorogenic, vanillic, and 4-hydroxy-3- methoxycinnamic acids showed a significant increase in concentration under W, with values exceeding 50% in most cases. Similarly, flavonoids such as myricetin and rutin exhibited the same behavior, whereas catechin was not affected by light treatment.

**Table 4.** Individual polyphenol concentrations in arugula, mustard, and radish microgreens grown under different LED light treatments.

| Polyphenol ($\mu\text{g g}^{-1}$ FW) | W | WB | R |
| --- | --- | --- | --- |
| **Arugula** |  |  |  |
| Galic acid | 0.91 ± 0.04 a | 0.43 ± 0.12 b | 0.55 ± 0.15 ab |
| 3,4-Dihydroxybenzoic acid | 0.26 ± 0.01 a | 0.03 ± 0.0006 b | 0.05 ± 0.011 ab |
| 4-Hydroxybenzoic acid | 3.03 ± 0.17 a | 2.01 ± 0.11 a | 2.27 ± 0.15 a |
| Ferulic acid | 5.32 ± 0.48 a | 4.61 ± 0.34 a | 4.41 ± 0.49 a |
| *p*-cumaric acid | 4.38 ± 0.33 a | 4.42 ± 0.46 a | 3.39 ± 0.26 a |
| Catechin | 6.36 ± 0.44 a | 6.41 ± 0.50 a | 6.06 ± 1.22 a |
| Myricetin | 1.67 ± 0.28 a | 1.42 ± 0.21 a | 1.08 ± 0.20 a |
| Quercetin-3-glucoside | 1.21 ± 0.19 a | 1.28 ± 0.11 a | 1.49 ± 0.22 a |
| **Mustard** |  |  |  |
| 3,4-Dihydroxybenzoic acid | 11.52 ± 1.43 a | 0.71 ± 0.35 b | 1.79 ± 0.32 b |
| 4-Hydroxybenzoic acid | 14.68 ± 1.67 a | 4.34 ± 0.65 b | 5.31 ± 0.52 b |
| Vanillic acid | 3.16 ± 0.33 a | 1.76 ± 0.18 b | 1.90 ± 0.05 b |
| Chlorogenic acid | 92.87 ± 14.24 a | 36.88 ± 2.04 b | 51.56 ± 6.01 b |
| Ferulic acid | 332.92 ± 19.38 a | 115.76 ± 11.43 b | 168.39 ± 11.05 b |
| *p*-cumaric acid | 1.19 ± 0.14 b | 1.92 ± 0.08 a | 1.51 ± 0.12 ab |
| Catechin | 71.05 ± 3.19 a | 85.14 ± 10.53 a | 77.07 ± 11.57 a |
| Myricetin | 30.87 ± 3.04 a | 7.65 ± 1.04 b | 14.87 ± 0.94 b |
| Rutin | 14.01 ± 1.64 a | 8.11 ± 0.49 b | 6.47 ± 0.87 b |
| **Radish** |  |  |  |
| 3,4-Dihydroxybenzoic acid | 0.18 ± 0.02 a | 0.10 ± 0.03 a | 0.21 ± 0.07 a |
| 4-Hydroxybenzoic acid | 9.28 ± 1.49 a | 5.47 ± 0.82 a | 6.08 ± 0.77 a |
| Vanillic acid | 12.25 ± 0.63 a | 14.55 ± 1.05 a | 12.30 ± 0.68 a |
| Chlorogenic acid | 19.96 ± 1.19 a | 19.73 ± 1.67 a | 19.65 ± 1.46 a |
| Ferulic acid | 109.95 ± 8.20 a | 112.86 ± 8.85 a | 99.40 ± 7.21 a |
| *p*-cumaric acid | 0.12 ± 0.002 a | 0.10 ± 0.02 ab | 0.05 ± 0.01b |
| Myricetin | 0.25 ± 0.04 a | 0.23 ± 0.004 a | 0.25 ± 0.03 a |
| Ellagic acid | 57.46 ± 9.38 a | 43.68 ± 4.61 a | 33.93 ± 3.33 a |

Values are expressed as mean ± standard error (n = 4). Different lowercase letters within each row indicate significant differences among light treatments within each species according to Tukey's test ( $P < 0.05$ ).

In contrast, p-coumaric acid showed an opposite response, reaching its highest concentration under treatment WB (1.92 μg g ¹ FW) compared to treatment W. The major compounds found in radish were 4-hydroxy-3-methoxycinnamic acid and ellagic acid, neither of which showed significant differences under the light treatments. The only phenol affected by spectral quality was p-coumaric acid, with its accumulation favored by treatment W (0.12 μg g ¹ FW) over treatment R (0.045 μg g ¹ FW). Overall, radish showed limited variation in the phenolic compounds identified under the different light treatments

### Integrated analysis of species responses to light treatments

In arugula, light treatments produced only minor changes across the evaluated traits, and no clear response pattern was observed among spectra (Figure 5). Most variables displayed similar standardized values regardless of the light treatment, indicating a limited effect of spectral composition on the overall physiological and biochemical profile of this species.

**Figure 5.**
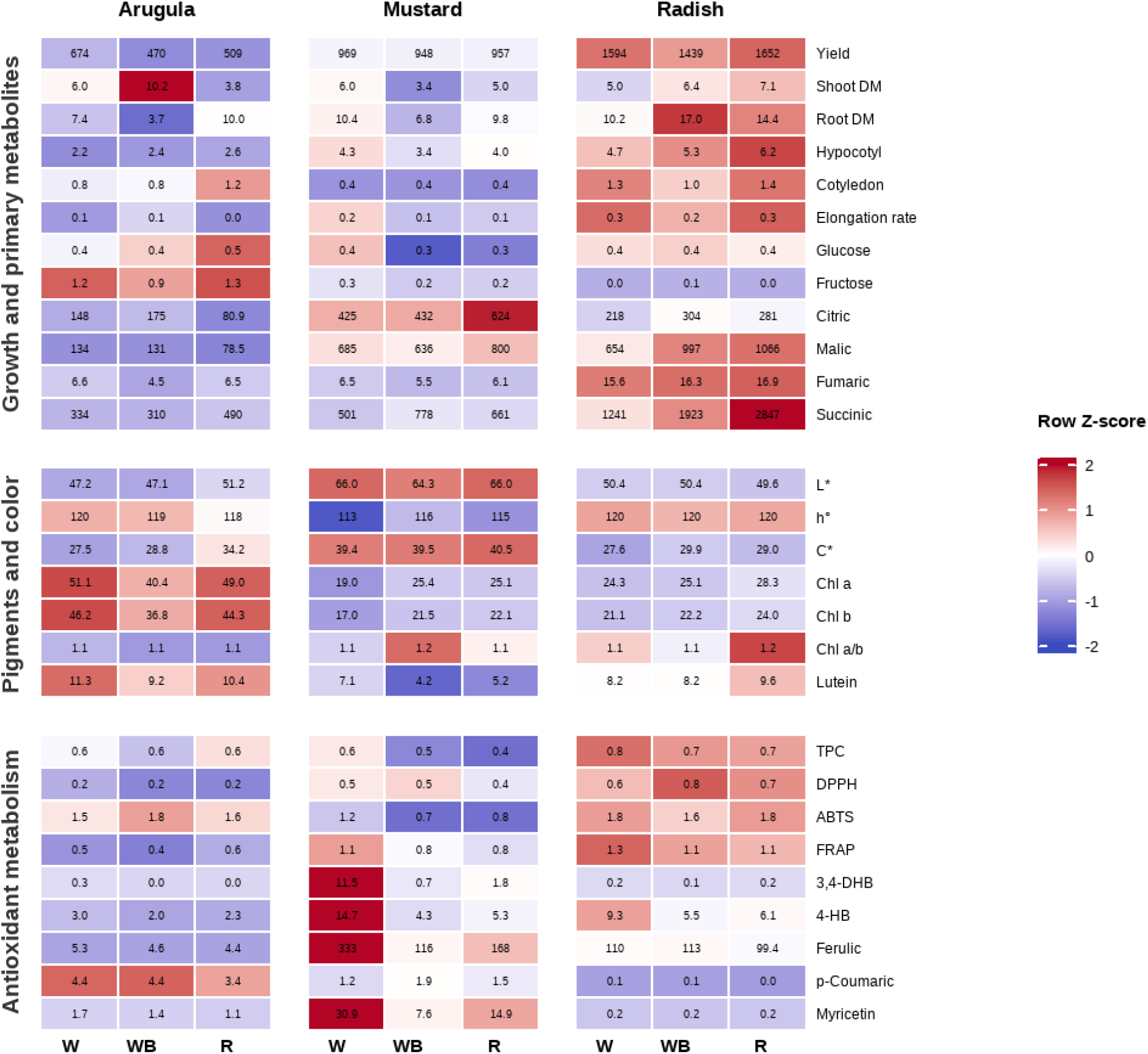
Heatmap of growth, biochemical, and antioxidant-related traits in arugula, mustard, and radish microgreens under different LED light treatments. Colors represent row-wise Z-scores. Values correspond to treatment means

Although the W treatment tended to exhibit higher standardized values for chlorophylls and lutein, no consistent trends were observed across the evaluated functional groups.

In mustard, the heatmap revealed only limited differences among light treatments, with no consistent response pattern across the evaluated functional groups. Most growth, pigment, and antioxidant-related variables displayed similar standardized values regardless of the spectral composition. The most evident effects were observed for citric acid, which reached its highest standardized values under the R treatment, and for some individual phenolic compounds, which were associated with the white spectrum (W).

In radish, the heatmap revealed a more distinct response to light treatments. Within the growth and primary metabolites block, the R treatment was frequently associated with the highest standardized values, whereas the antioxidant metabolism block showed no consistent response pattern among light treatments. These results indicate that the clearest responses to spectral quality were observed in growth and primary metabolism, whereas antioxidant-related variables exhibited comparatively smaller and less consistent changes under the conditions of this study.

## Discussion

The results of this study showed that, although the three species evaluated belong to the Brassicaceae family, they exhibited distinct responses to the light spectra applied, in agreement with previous reports (Flores et al., 2024; Toscano et al., 2021; Ying et al., 2020a).

The spectral composition influenced morphological traits and dry matter accumulation, depending on the species evaluated, but this was not reflected in biomass yield, which was independent of the spectrum used (Table 2).

The results obtained regarding dry matter content in both the shoot and root suggest species-specific differences in tissue water content and, possibly, biomass allocation. The arugula showed the highest shoot dry matter content under treatment WB, in contrast to treatment R, which, although it promoted greater elongation, had the lowest shoot dry matter content. This is consistent with the finding of Toscano et al. (2021), who obtained the lowest dry matter content in turnip and amaranth under red light.

In contrast, the highest shoot dry matter values were observed under R in radish and under W in mustard, while WB showed intermediate values in radish. Thus, increased hypocotyl elongation was not accompanied by a reduction in shoot dry matter accumulation. While the low proportion of blue light may have promoted elongation responses associated with reduced blue-light perception, it did not negatively impact dry biomass accumulation. Others have reported increased dry matter in green and red kale under white-blue and white-red light compared with white light (Frąszczak et al., 2023). However, the statistical similarity observed between treatments R and W in mustard and R and WB in radish reinforces the idea of plasticity in these species.

Regarding root dry matter, arugula showed the highest dry matter content under treatment R, with no differences compared to treatment W. This response may reflect differences in biomass allocation and/or tissue hydration patterns. Conversely, mustard and radish showed no differences in root dry matter content under the light treatments. In lettuce, higher root dry and fresh weights were reported when grown under light with peaks in the red and blue, and a spectral band of 500-600 nm, compared to monochromatic red and blue light (Lin et al., 2013).

Treatment R promoted the greatest hypocotyl elongation in arugula and radish, with no differences compared to treatment WB. Although treatment R does not consist of monochromatic red light, the low proportion of blue light (4%) may have been insufficient to induce cryptochrome-mediated inhibition of hypocotyl elongation (Huché-Thélier et al., 2016).

In mustard, however, the response differed from that observed in arugula and radish. Seedlings grown under W exhibited the greatest hypocotyl length, whereas WB resulted in the shortest values. This may suggest a higher sensitivity of mustard to blue light-mediated inhibition of hypocotyl elongation. Under WB, the 15% proportion of blue light may have been sufficient to suppress elongation, whereas under W the lower proportion of blue light (4%) did not produce the same response.

In a study by Ying et al. (2020b), hypocotyl elongation in arugula and cabbage was not affected by up to a 30% increase in blue light; however, mustard and kale showed reduced hypocotyl elongation at higher proportions of blue light. In contrast, when evaluating the hypocotyl elongation rate, no clear relationship was observed between final hypocotyl length and elongation rate across species. Arugula showed the lowest elongation rate under treatment R, despite its greater final length. These differences could be attributed to the timing of the measurements, omitting the growth that occurred in earlier stages. It is worth noting that the microgreens remained in darkness for 4 days, a period during which the elongation rate could have been higher due to etiolation (Li et al., 2023b). This stage could also explain the lack of differences in the radish elongation rate.

Furthermore, the effect of light quality on cotyledon area differed among species. In one study, basil, mustard, and pea microgreens showed larger cotyledon areas when grown under light with a higher red/blue ratio. In contrast, radish, chives, and borage showed no differences across light treatments (Bantis et al., 2021). Others found that arugula, kale, mustard, and cabbage had larger cotyledon areas under monochromatic red light and the smallest values under blue and green light (6%) (Kong & Zheng, 2020). Similarly, in this study, treatment R promoted maximum cotyledon expansion in arugula. Conversely, radish and mustard showed no significant differences between light treatments. Plant morphological responses to light conditions differ among species because they exhibit varying degrees of phenotypic plasticity (Kong & Zheng, 2018).

In this study, color variables were differentially affected by the light spectrum across species. Mustard showed higher lightness values under W and R, accompanied by a lower hue angle under W. This is consistent with a more yellow coloration under this spectrum. Similarly, arugula showed higher L* and C* values but a lower hab under R. Color parameters are directly related to plant pigments, with chlorophylls and carotenoids being the main pigments affecting their appearance (Vrkić et al., 2024). However, despite these changes in color, no differences were found in chl a and b content or total carotenoids across any species or treatments evaluated.

The effects of LED spectral quality on photosynthetic pigment accumulation remain inconsistent across studies, suggesting that no universal response to individual wavelengths exists. Craver et al. (2017) observed that the effects of light quality differed among Brassica species and proposed that the accumulation of photosynthetic pigments exhibited species-dependent responses. Similarly, Toscano et al. (2021) found that blue light increased chlorophyll accumulation in amaranth microgreens but had no significant effect in turnip greens grown under identical conditions. Hernández-Adasme et al. (2026) introduced an additional level of complexity by demonstrating that chlorophyll responses were influenced by interactions among light spectrum, irradiance, and genotype. Taken together, these findings indicate that the regulation of photosynthetic pigments under artificial lighting depends on both the spectral environment and species-specific physiological characteristics. Although Chl a, Chl b, and total chlorophyll remained largely unchanged in the present study, the Chl a/b ratio was affected in mustard and radish, suggesting subtle modifications in the organization of the photosynthetic apparatus rather than changes in the overall chlorophyll pool. The Chl a/b ratio is widely recognized as an indicator of acclimation to different light environments because Chl b is predominantly associated with the light-harvesting antenna complexes. In contrast, Chl a is more abundant in the reaction centers of photosystems. Consequently, changes in this ratio may reflect modifications in the relative size or composition of the light-harvesting complexes that optimize light capture under different spectral conditions. Similar interpretations have been proposed for plants exposed to shade or modified light environments, where decreases in the Chl a/b ratio have been associated with adjustments in light-harvesting organization rather than changes in photosynthetic capacity (Shao et al., 2014; Wang & Folta, 2013).

Lutein, one of the major xanthophylls in green tissues, plays important roles in light harvesting and photoprotection by stabilizing the photosynthetic apparatus and dissipating excess excitation energy. Consequently, changes in lutein concentration under different light environments have often been interpreted as part of the plant acclimation response. However, the effects of LED spectral quality on lutein accumulation remain inconsistent across studies. While several authors have reported higher lutein concentrations under blue-enriched light conditions (Lee et al., 2023; Samuolienė et al., 2017), Kopsell et al. (2014) found no significant differences among LED spectral combinations in broccoli microsprouts. Likewise, Craver et al. (2017) highlighted that carotenoid responses to light quality are highly species-dependent and suggested that the spectral differences applied may not always be sufficient to induce significant changes in pigment accumulation.

In the present study, lutein concentration was affected only in mustard, whereas no significant changes were detected in arugula or radish, despite all three species being grown under identical environmental conditions. This species-specific response is consistent with previous reports indicating that lutein accumulation under different light environments varies among species.

The response of soluble sugars to light treatments was species-dependent. In arugula and radish, no significant differences in glucose and fructose content were observed among treatments (Figure 3). In mustard, treatment W showed higher glucose and fructose contents than WB and R. These results suggest that soluble sugar accumulation in mustard was more responsive to spectral composition than in the other species evaluated.

This partner is consistent with previous studies reporting greater accumulation of both sugars in mustard and amaranth when grown under 88.9% red and 11.1% blue light (Gudžinskaitė et al., 2025). Similarly, broccoli sprouts showed increased fructose content under white, blue, and violet light, while glucose increased under these spectra as well as red, green, and yellow light (Zhuang et al., 2022). Likewise, increased glucose levels have been reported in basil and dill under broad-spectrum red light (El Haddaji et al., 2023).

In this study, spectral quality modified the content of organic acids in a species-dependent manner. In arugula, the content of citric and malic acids decreased in response to treatment R. A similar behavior was observed in spinach when treated with a spectrum containing 74% red light at a light intensity of 260 μmol m ² s ¹ (Hernández-Lara et al., 2026). Treatment R increased succinic acid accumulation in both arugula and radish. Succinic acid was also the predominant organic acid in both species. According to Fedorin et al. (2022), some enzymes of the Krebs cycle are light-sensitive; in particular, succinate dehydrogenase decreases its activity under red light. Therefore, the increase in succinic acid could be explained by the lower activity of this enzyme.

In mustard, however, malic acid was the predominant acid, with no significant differences between the treatments studied. Contrary to arugula and radish, succinic acid showed greater accumulation under treatments W and WB. Fumaric acid, on the other hand, did not show significant differences between the light treatments for the three species.

Ascorbic acid content in radish was affected by the light spectrum, increasing significantly under treatment R. In mustard, light treatments had no effect on ascorbic acid content. In amaranth and turnip microgreens, higher ascorbic acid content was found under monochromatic blue light with a 16 h photoperiod (Toscano et al., 2021). In broccoli, the greatest increase in AA content was observed under a spectrum composed of blue, red, and green light (Samuoliene et al., 2019). This suggests that AA regulation depends on both spectral composition and species.

In arugula, treatment R resulted in higher TAC when determined by the FRAP method, whereas TPC did not differ among the spectra (Figure 4). This suggests that polyphenols were probably not the main contributors to the observed variation in antioxidant capacity. Although red light has been reported to increase vitamin C and certain phenolic acids in some species (Lee et al., 2019; Samuolienė et al., 2016), opposite responses have also been described (Ali et al., 2025). Under the conditions of this study, neither ascorbic acid nor the identified phenolic compounds increased under treatment R, suggesting that the quantified phenolic compounds alone do not fully explain the observed antioxidant capacity.

Mustard showed higher TAC under treatment W when evaluated by the ABTS method. Since TPC and most of the identified phenolic compounds also increased under this treatment, the higher antioxidant capacity was probably associated with the accumulation of phenolic compounds. In radish, TAC measured by the DPPH assay was higher under treatment WB, whereas TPC remained unchanged among treatments. This suggests that the observed differences in antioxidant capacity were associated with changes in specific antioxidant compounds rather than total phenolic content. However, DPPH measurements may be influenced by interference from anthocyanins (Gülçin, 2020).

In arugula, spectral quality had a limited effect on the composition of individual phenolic compounds, with significant variation observed only in 3,4-dihydroxybenzoic and gallic acids. Similarly, the phenolic profile of radish remained largely unaffected by the light treatments, with significant differences detected only for p-coumaric acid. This limited response agrees with previous reports showing that genotype plays a dominant role in determining the phenolic profile of radish regardless of growing conditions (Silva et al., 2025; Mlinarić et al., 2023). The higher accumulation of p-coumaric acid under treatment W was also consistent with the findings of Silva et al. (2025), whereas the remaining phenolic compounds showed no consistent response to spectral quality.

Unlike arugula and radish, mustard microgreens exhibited a marked response to spectral quality. Consistent with the increases observed in TPC and TAC, most of the identified phenolic compounds accumulated to higher concentrations under W treatment. Hydroxycinnamic acids constituted the predominant phenolic fraction, in agreement with previous reports for Brassica microgreens (Kyriacou et al., 2019; Oszmiański et al., 2013).

These results suggest that, in mustard, spectral quality promoted a broader activation of phenolic metabolism than in the other species.

## CONCLUSION

The response to light quality was strongly species-dependent. In radish, light treatments affected only a limited number of traits, whereas in arugula significant effects were observed mainly in morphology and selected biochemical traits. In mustard, light treatments significantly affected sugars, phenolic composition, total phenolic content, and antioxidant capacity. These findings demonstrate that responses to LED spectral composition cannot be generalized across Brassicaceae microgreens, as the optimal light treatment depended on both the species and the quality attribute of interest. Therefore, light management should be optimized according to the target species and the desired quality attributes.

---

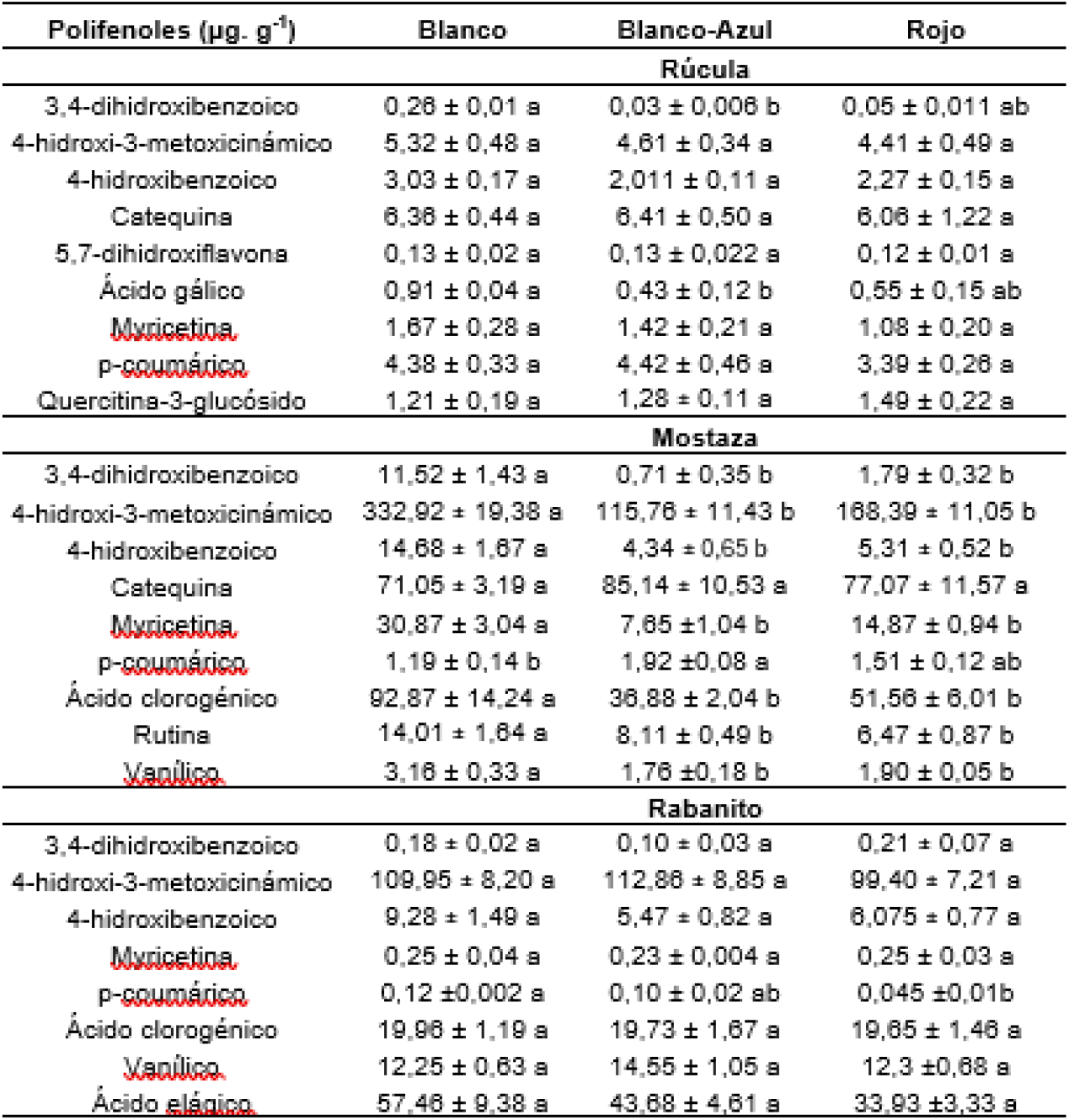

